## Supplementary data for "Molecular determinants of inhibition of UCP1-mediated respiratory uncoupling"

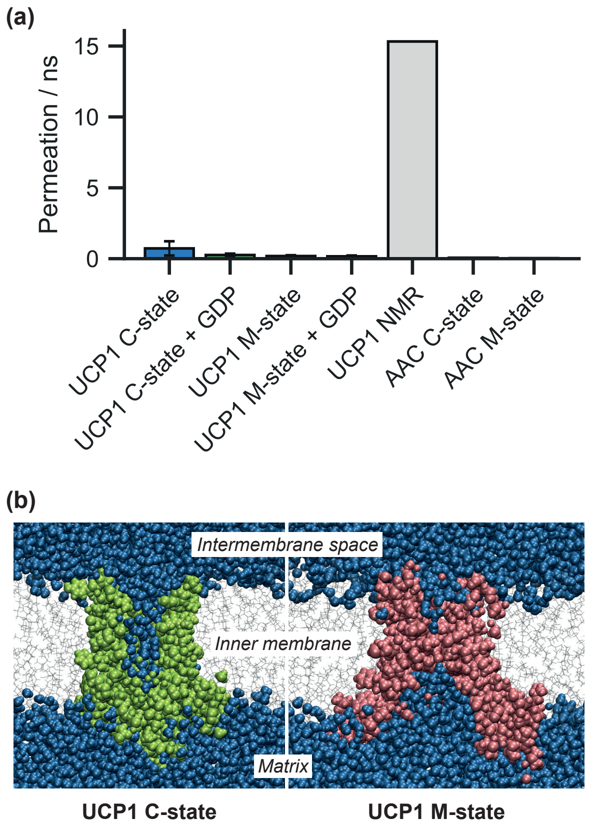


Figure S1: **Water permeability measurements in MD simulations.** **(a)** Number of water molecules crossing the membrane through the protein per nanosecond. UCP1 NMR is an homology model of UCP1 built from NMR structure of UCP2 (PDB 2LCK). Error bar is the Standard Error of the Mean between replicas averages. **(b)** Cross-section view of UCP1 hydration. Water molecules are depicted as blue spheres and atoms of UCP1 C-state and UCP1 M-state are represented as, respectively, green and red spheres.

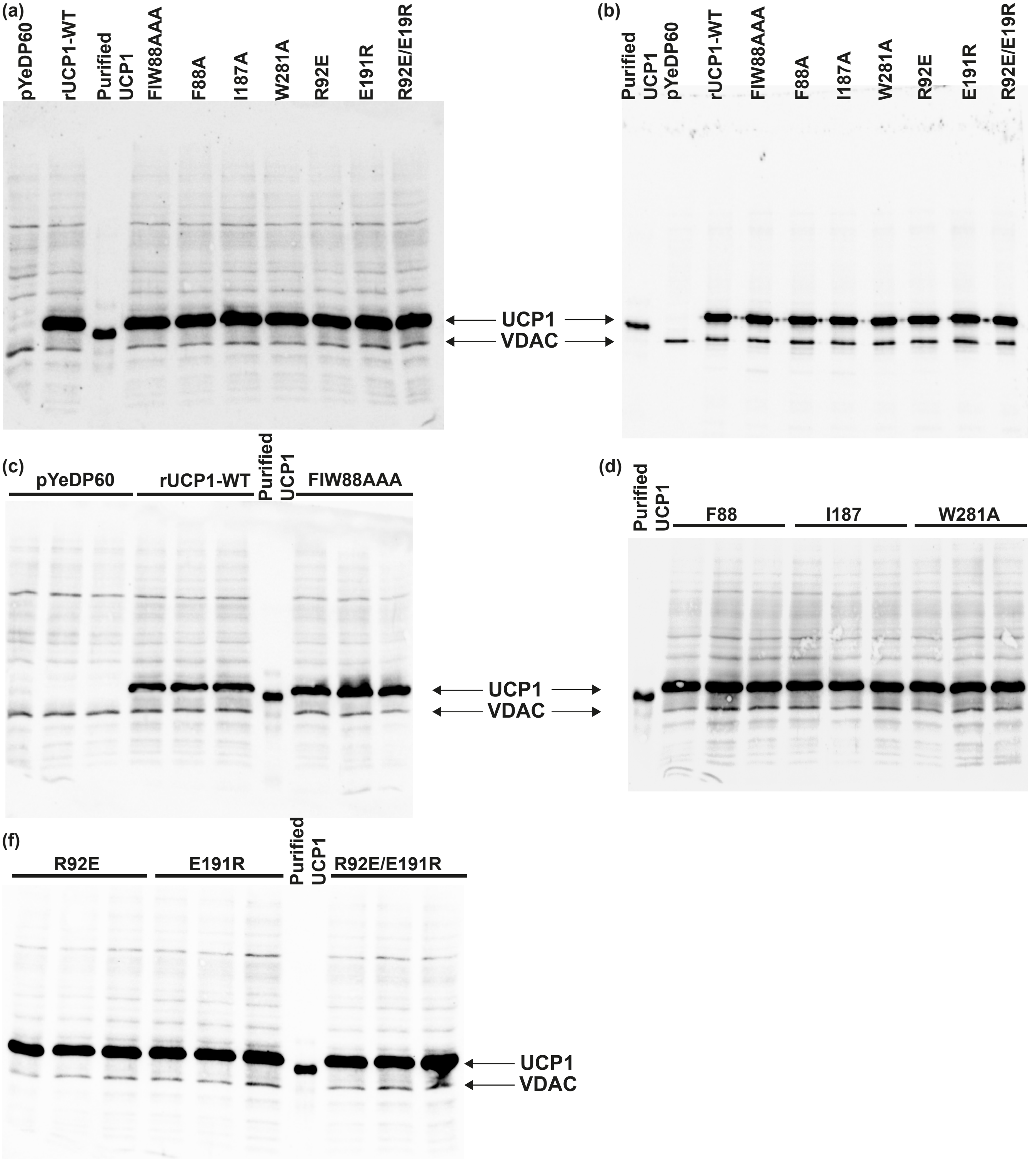


Figure S2: **Immunodetection of UCP1 on recombinant yeast grown in S-lactate medium and prepared according to Methods.** Immunodetection of VDAC is used as a loading control. Both proteins are revealed using a mouse anti-pentahistine tag:HRP and a mouse anti-VDAC1, see Methods. Expression is measured on total TCA extracts (a) and on mitochondria (b). (c),(d),(e) are replicates of total TCA extracts. Statistical analyses are presented in Table S1.

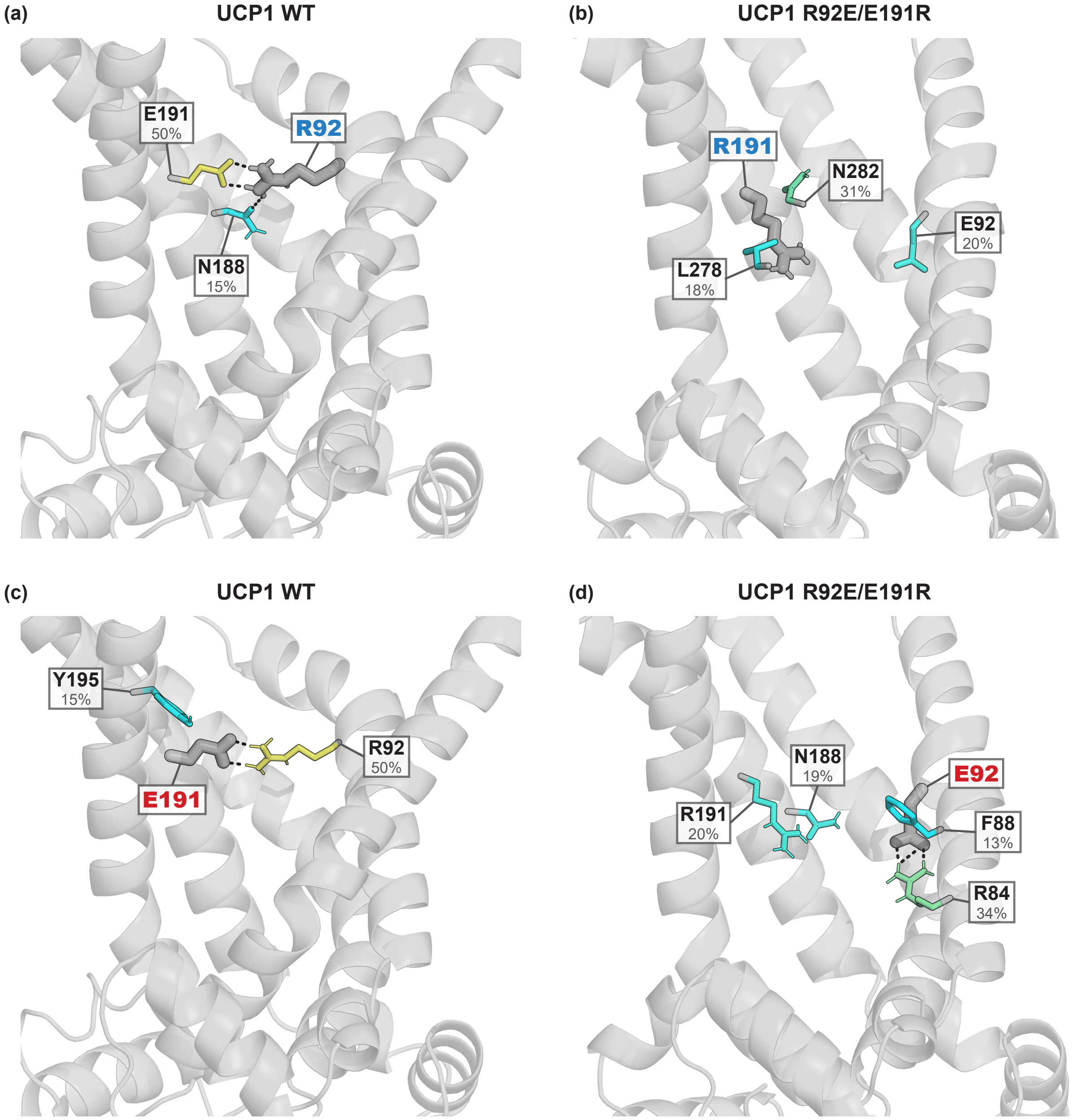


Figure S3: **Intra-protein contacts in simulations of UCP1 wild type and the double mutant R92E/E191R.** **(a)**, **(b)**, **(c)** and **(d)** Contact time between protein residues and the charged atoms of, respectively, R92, R191, E191 and E92. Snapshots of trajectories at 500 ns of UCP1 wild type for **(a)** and **(c)**, and of UCP1 R92E-E191R for **(b)** and **(d)**. Only the residues with a contact time higher than 10% are represented. Residues are colored according to their contact time. The color scale goes from cyan at 10% to yellow at 55% to red at 100%. Contact times are computed on simulations from 200 ns to 1000 ns.

Figure S4: **Multiple alignments of human mitochondrial carrier sequences and rat UCP1.** Gray bars are indicate hidden parts of sequences. Colored backgrounds are an estimation of alpha helices from the UCP1 C-state model gray and yellow backgrounds are respectively odd and even transmembrane helices, and green are non-transmembrane helices. Dark gray vertical bars depict hidden parts of the sequences. Abbreviations: *rUCP1* rat Uncoupling Protein 1; *hsUCP1* Uncoupling Protein 1; *hsUCP2* Uncoupling Protein 2; *hsUCP3* Uncoupling Protein 3; *hsUCP4* Uncoupling Protein 4; *hsUCP5* Uncoupling Protein 5; *hsDIC* Mitochondrial dicarboxylate carrier; *hsODC* Mitochondrial 2-oxodicarboxylate carrier; *hsTXTP* Tricarboxylate transport protein; *hsAAC1* ADP/ATP Carrier 1; *hsGDC* Graves disease carrier; *hsSCMC1* Mitochondrial ATP-Mg/Pi carrier protein 1; *hsS2536* Solute carrier family 25 member 36; *hsMFTC* Mitochondrial folate transporter/carrier; *hsS2540* Solute carrier family 25 member 40; *hsS2538* Mitochondrial glycine transporter; *hsCMC1* Mitochondrial aspartate glutamate carrier 1; *hsMPCP* Phosphate carrier protein.

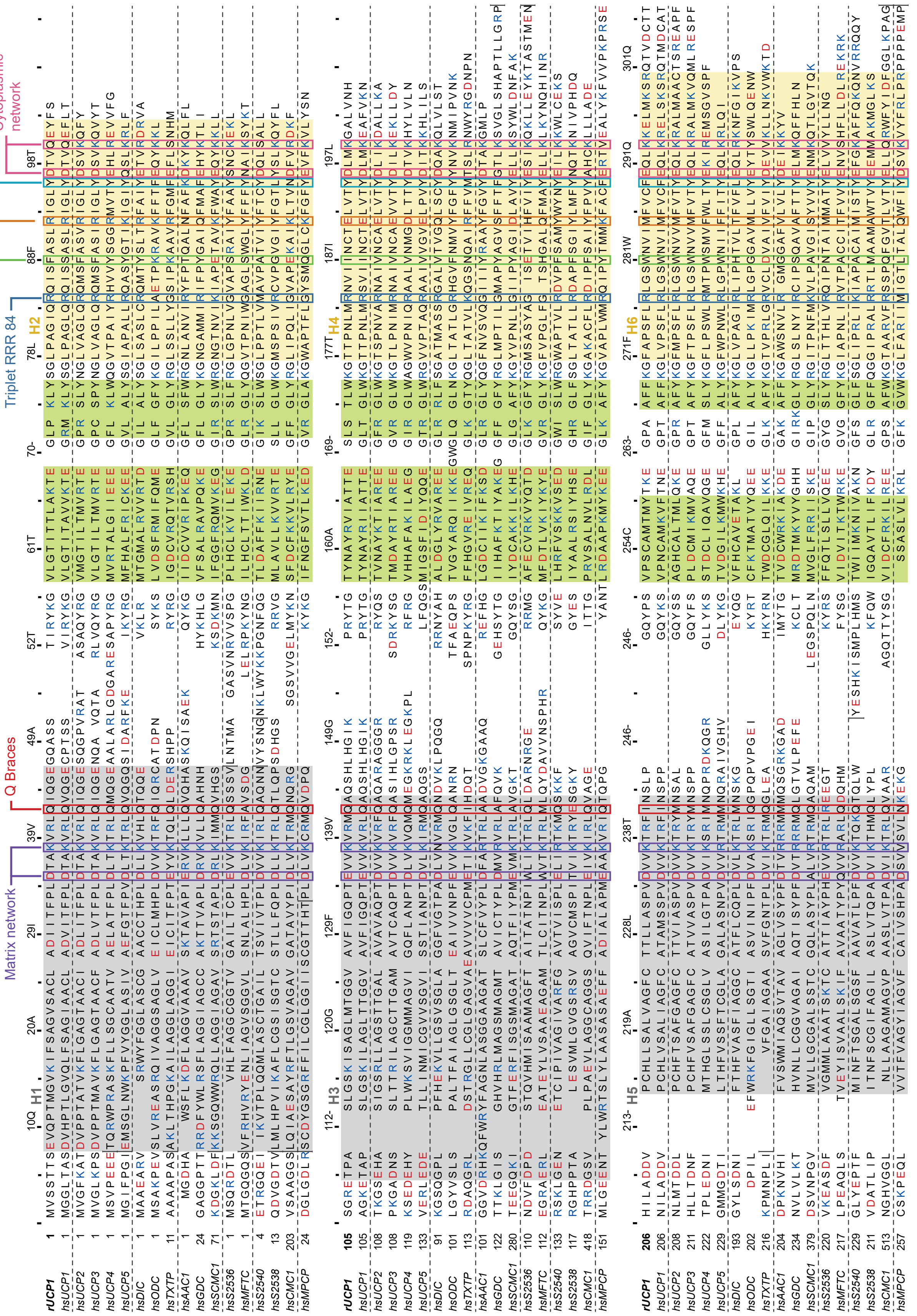


Legend on previous page.

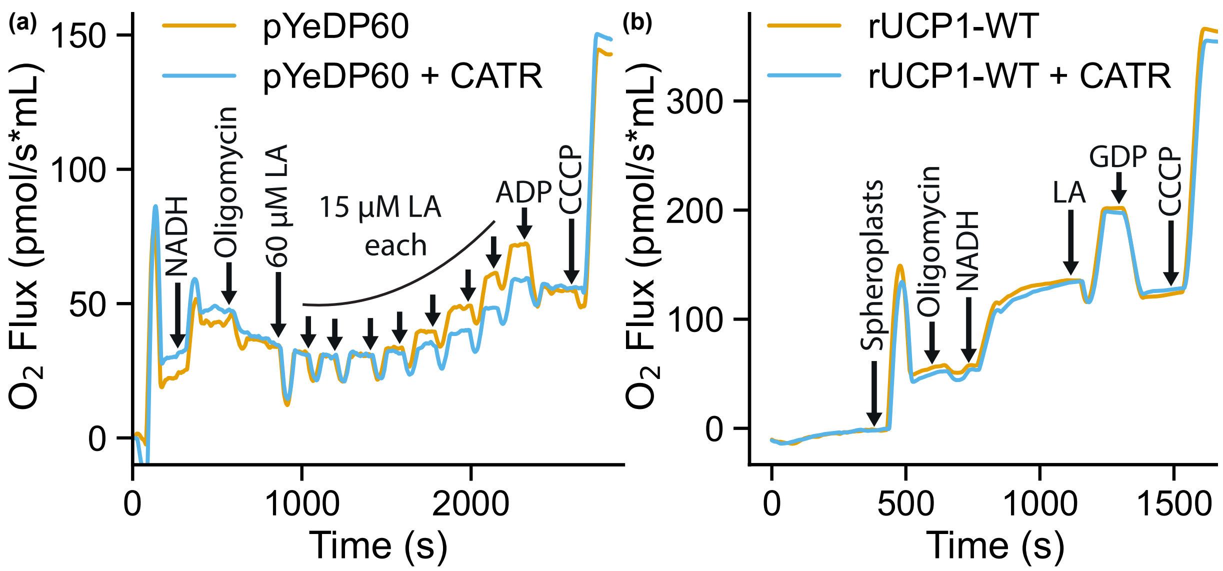


Figure S5: **AAC-dependent respiratory uncoupling is inhibited by ADP and requires a higher FFA concentration than UCP1-dependent uncoupling.** **(a)** Respiration curves showing the effect of 60 micromolar concentration of LA and of subsequent multiple additions of 15 micromolar amount of FFA to yeast control spheroplasts in the presence (blue curve) or absence (orange curve) of the CATR inhibitor of AAC. As shown in [15], the addition of ADP suppresses, as well as CATR, the LA-induced increase of respiration. **(b)** Respiration curves of rUCP1-WT spheroplasts in the presence (blue curve) or absence (orange curve) of CATR. In this experimental setup, the addition of 60 micromolar concentration of LA (LA/BSA=4) had no effect on yeast AAC.

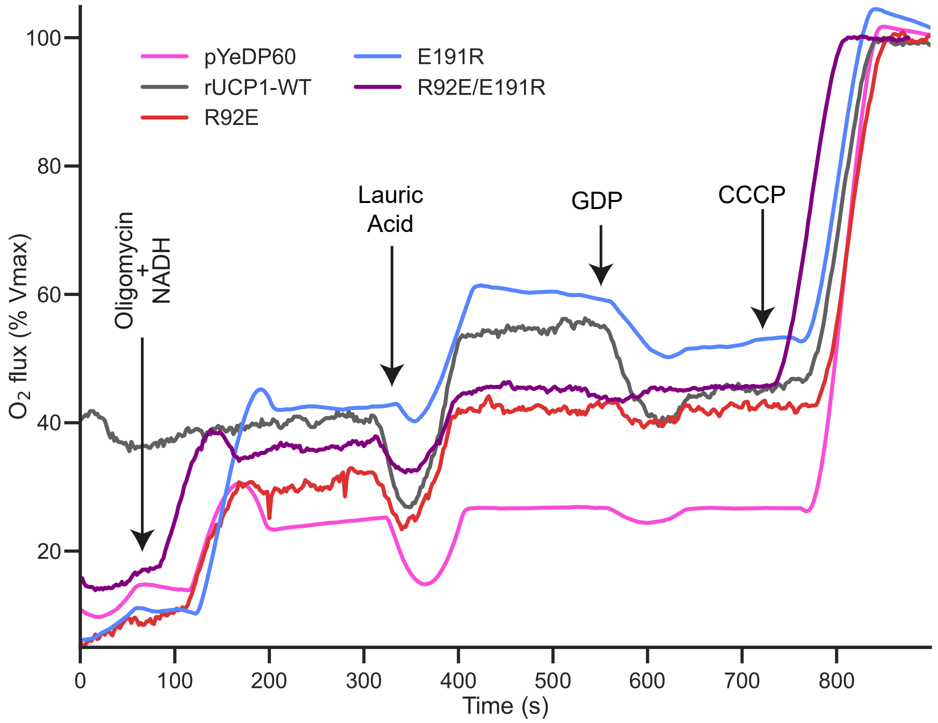


Figure S6: **Representative oxygen flux consumption curves of permeabilized spheroplasts harboring either control pYeDP60 plasmid expression (pink curve) or expressing rUCP1-WT (grey curve), mutants R92E (red curve), E191R (blue curve), R92E/E191R (purple curve).**

| Mutant | Summary | P value[1] | Mean rUCP1 | Mean mutant | Mean diff |
|---|---|---|---|---|---|
| R92E | ns | 0.1221 | 1.859 | 3.120 | 1.260 |
| E191R | ns | 0.9447 | | 2.291 | 0.4318 |
| R92E/E191R | ns | 0.9995 | | 2.049 | 0.1893 |
| FIW88AAA | ns | 0.9770 | | 2.221 | 0.3618 |
| F88A | ns | > 0.9999 | | 1.838 | -0.021 |
| I187A | ns | 0.9996 | | 1.725 | -0.1342 |
| W281A | ns | > 0.9999 | | 1.829 | -0.03 |

Table S1: **Statistical analyses of UCP1 wild-type and mutant expression in yeast total TCA extracts.** UCP1/VDAC ratios are calculated using the SDS-PAGES shown in Figure S2. ImageJ 1.53k and statistical analyses calculate intensity, are performed on PRISM with One-way ANOVA and Dunnett's multiple comparison test with rUCP1-WT. ns = non significant, Standard Error of difference = 0.5412, Degrees of freedom = 32, F value = 1.397, N = 6. [1]: P value adjusted for multiple tests.

| Mutant | Summary | P value[1] | Mean rUCP1-WT | Mean mutant | Différence Moyenne |
|---|---|---|---|---|---|
| pYeDP60 | ** | 0,0042 | 1,667 | 2,512 | -0,8448 |
| R92E | ns | 0,9999 | | 1,662 | 0,004974 |
| E191R | ns | 0,9929 | | 1,473 | 0,1937 |
| R92E/E191R | ns | 0,9999 | | 1,604 | 0,06268 |
| FIW88AAA | ns | 0,9134 | | 1,371 | 0,2961 |
| F88A | ns | 0,9999 | | 1,746 | -0,07914 |
| I187A | ns | 0,8510 | | 1,335 | 0,3318 |
| W281A | ns | 0,9865 | | 1,453 | 0,2139 |

Table S2: **Statistical analyses of spheroplasts respiratory control ratio (RCR) with One-way ANOVA and Dunnett's multiple comparison test versus rUCP1-WT.** Standard Error of difference: pYeDP60 = 0.2112, rUCP1-WT and mutants = 0.2587, Degrees of freedom = 24, F value = 4.704, N: pYeDP60 = 6, rUCP1-WT and mutants = 3. [1]: P value adjusted for multiple tests.

| Mutant | Summary | P value[1] | Mean rUCP1 | Mean mutant | Mean diff |
|---|---|---|---|---|---|
| R92E | ** | 0.0441 | 27.26 | 19.37 | 7.889 |
| E191R | ns | 0.1140 | | 20.77 | 6.486 |
| R92E/E191R | * | 0.01 | | 17.44 | 9.818 |

Table S3: **Statistical analyses of spheroplasts respiration activation by lauric acid with One-way ANOVA and Dunnett's multiple comparison test: rUCP1-WT versus salt bridge mutants.** Standard Error of difference = 3.173, Degrees of freedom = 37, F value = 3.433, N = 11. [1]: P value adjusted for multiple tests.

| Mutant | Summary | P value[1] | Mean rUCP1 | Mean mutant | Mean diff |
|---|---|---|---|---|---|
| R92E | **** | < 0.0001 | 98.11 | -16.18 | 114.3 |
| E191R | **** | < 0.0001 | | 32.91 | 65.20 |
| R92E/E191R | *** | < 0.0001 | | -13.45 | 111.6 |

Table S4: **Statistical analyses of UCP1 inhibition by GDP with One-way ANOVA and Dunnett's multiple comparison test: rUCP1-WT versus salt bridge mutants.** Standard Error of difference = 11.67, Degrees of freedom = 38, F value = 41.18, N = 11. [1]: P value adjusted for multiple tests.

| Mutant | Summary | P value[1] | Mean rUCP1 | Mean mutant | Mean diff |
|---|---|---|---|---|---|
| FIW88AAA | **** | < 0,0001 | 48.11 | 81.03 | -31.66 |
| F88A | ns | 0,9794 | | 43.76 | 5.615 |
| I187A | **** | < 0,0001 | | 75.83 | -26.46 |
| W281A | **** | 0.0002 | | 74.05 | -24.67 |

Table S5: **Statistical analyses of spheroplasts respiration activation by lauric acid with One-way ANOVA and Dunnett's multiple comparison test: rUCP1-WT versus 88 mutants.** Standard Error of difference = 5.546, Degrees of freedom = 50, F value = 18.72, N = 11. [1]: P value adjusted for multiple tests.

| Mutant | Summary | P value[1] | Mean rUCP1 | Mean mutant | Mean diff |
|---|---|---|---|---|---|
| FIW88AAA | **** | <0.0001 | 87.77 | -4.536 | 92.31 |
| F88A | ns | 0.7064 | | 94.30 | -6.524 |
| I187A | **** | <0.0001 | | 24.09 | 63.68 |
| W281A | **** | <0.0001 | | 20.12 | 67.66 |

Table S6: **Statistical analyses of UCP1 inhibition by GDP with One-way ANOVA and Dunnett's multiple comparison test: rUCP1-WT versus 88 mutants.** Standard Error of difference = 6.466, Degrees of freedom = 50, F value = 92.89, N = 11. [1]: P value adjusted for multiple tests.

| Addition | UCP1 | Summary | P-value[1] | Mean control | Mean UCP1 | Mean diff | SE of diff | F value |
|---|---|---|---|---|---|---|---|---|
| GDP | **rUCP1-WT** | **** | < 0,0001 | 0,2631 | -18,29 | 18,56 | 2.65 | 47.62 |
| | **FIW88AAA** | ns | 0,7199 | | 1,989 | -1,726 | | |
| AL 1 | **rUCP1-WT** | * | 0,025 | 2,089 | -11,03 | 13,12 | 4.973 | 147.2 |
| | **FIW88AAA** | **** | < 0,0001 | | 57,89 | -55,8 | | |
| AL 2 | **rUCP1-WT** | ns | 0,7873 | 8,572 | 12,00 | -3,431 | 6.235 | 132.2 |
| | **FIW88AAA** | **** | < 0,0001 | | 88,04 | -79,47 | | |

Table S7: **Statistical analyses of GDP addition then lauric acid on UCP1.** Analyses are done with One-way ANOVA and Dunnett's multiple comparison test with pYeDP60. Degrees of freedom = 25, N = 11. [1]: P value adjusted for multiple tests.

Video S1: **GDP entry into the UCP1 common substrate binding site, as observed in an ABMD simulation (simulated time shown: 20 ns), and rendered using VMD.** The R92/E191 salt bridge and arginine triplet R84 are represented in licorice, colored in orange and cyan respectively. GDP is colored by element.
